# Higher Magnetic Field NMR Renders Resolution Enhancement on Ganglioside GD3 Catalyzed Aβ_42_ Aggregates

**DOI:** 10.64898/2026.03.15.711644

**Authors:** Jhinuk Saha, Thirupati Ravula, Ayyalusamy Ramamoorthy

## Abstract

Magic-angle spinning (MAS) solid-state NMR (SSNMR) has been widely used to determine amyloid fibril structures at atomic resolution. Such studies typically rely on homogeneous fibril preparations that produce narrow linewidths and high spectral resolution, enabling reliable resonance assignment and structural analysis. However, many biologically relevant amyloid aggregates are structurally heterogeneous, resulting in spectral broadening and reduced sensitivity that hinder atomic-resolution characterization. Lipids are known to modulate amyloid aggregation pathways and promote the formation of toxic species that are often less homogeneous, further complicating NMR-based investigations. Here, we evaluate the feasibility of utilizing the benefits associated with high-field (1.1 GHz) SSNMR for studying ganglioside GD3–catalyzed Aβ_42_ aggregates. Uniformly-^13^C,^15^N-labeled Aβ_42_ was incubated with GD3 to generate lipid-associated aggregates and analyzed under MAS conditions. ^13^C cross-polarization magic-angle spinning (CPMAS) spectra and 2D ^13^C-^13^C chemical shift correlation experiments using CORD (*CO*mbined *R*2_*n*_^*v*^-*D*riven) mixing were acquired and compared with data collected at 600 MHz. Despite the heterogeneous nature of the GM1-associated assemblies, the 1.1 GHz spectra exhibit enhanced sensitivity and improved spectral resolution. Better resolved resonances corresponding to selectively structured regions of Aβ_42_ are observed, indicating the presence of an ordered core within the lipid-associated aggregates. These results demonstrate that ultrahigh-field SSNMR significantly improves the characterization of heterogeneous amyloid assemblies and provides a promising approach for atomic-level investigation of biologically relevant, lipid-modulated Aβ aggregates.

## Introduction

Self-assembly and aggregation of amyloid-β (Aβ) peptides (Fig.1a) are central events in the pathology of Alzheimer’s disease (AD).^1^ The conversion of soluble Aβ monomers into oligomers, protofibrils, and mature fibrils is associated with neurotoxicity and disease progression.^2–4^ Structural characterization of these aggregated states is therefore essential for understanding the molecular mechanisms underlying AD and for guiding therapeutic development. Over the past two decades, solid-state NMR (SSNMR) spectroscopy has emerged as one of the most powerful techniques for determining the structure and dynamics of amyloid fibrils at atomic resolution.^5–8^ In particular, magic-angle spinning (MAS) based SSNMR experiments enable site-specific resonance assignment, secondary structure determination, and the identification of intermolecular contacts in non-crystalline protein assemblies.^9–11^

**Figure 1.**
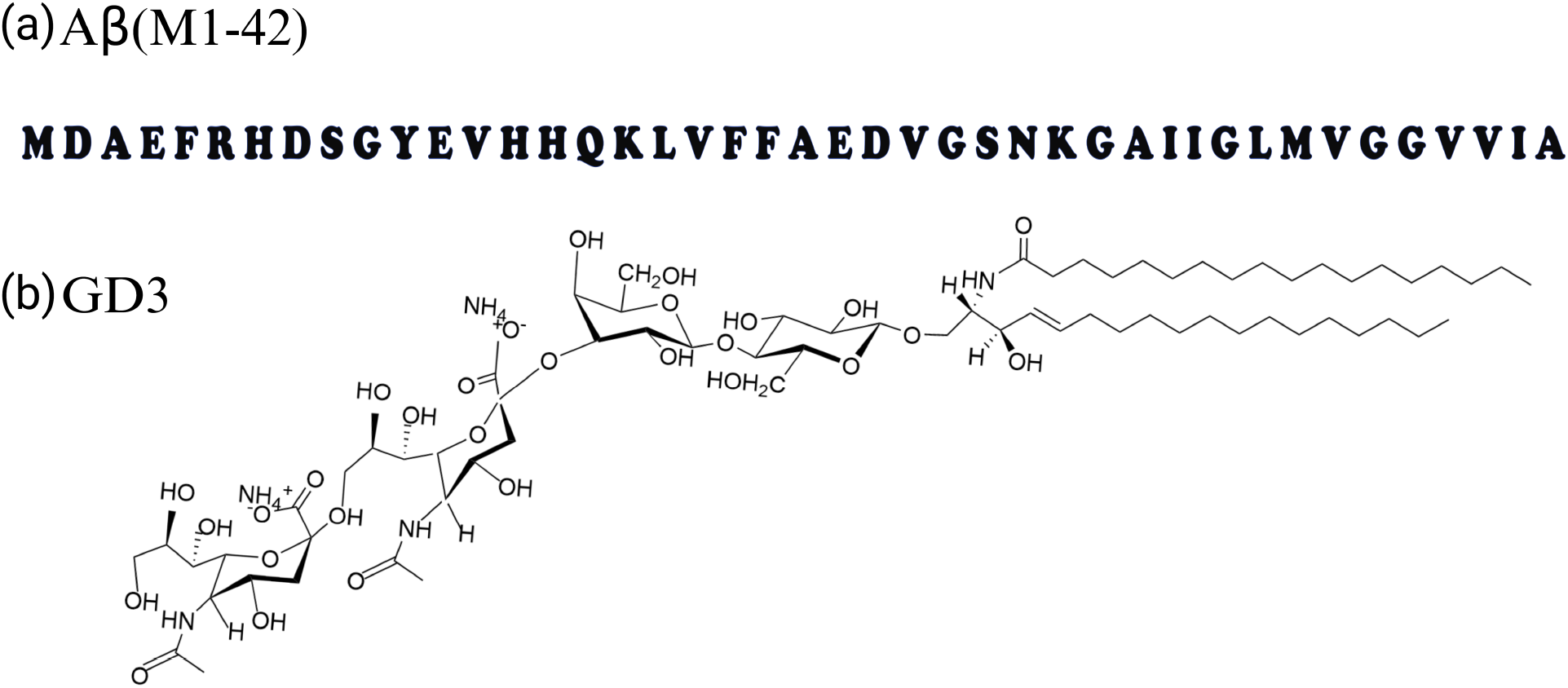
(a) Amino acid sequence of Aβ(M1-42) and (b) molecular structure of ganglioside, GD3, used in this study.

High-resolution SSNMR studies typically rely on structurally homogeneous fibril preparations, which yield narrow spectral linewidths and well-resolved resonances that facilitate assignment and structural analysis.^6,11,12^ However, amyloid aggregates formed under physiologically relevant or lipid-associated conditions are often structurally heterogeneous. Such heterogeneity results in spectral broadening, reduced sensitivity, and overlapping resonances, thereby complicating atomic-resolution structural studies.^13–16^ These challenges are especially pronounced for lipid-associated Aβ assemblies, which are increasingly recognized as biologically relevant and potentially more toxic species.^17–22^ Lipids are known to play critical roles in AD pathology^23–27^. Gangliosides, in particular, have been shown to interact with Aβ peptides (Fig.1a), modulate their conformational ensemble, and catalyze aggregation.^28–32^ Among them, ganglioside GD3 (Fig.1b) is abundant in neuronal membranes and may influence Aβ aggregation pathways and aggregate morphology^33–36^. Despite growing evidence for lipid-mediated modulation of Aβ aggregation, detailed structural information on such heterogeneous, lipid-associated assemblies remains limited.

In this study, we investigate the effect of ganglioside GD3 on Aβ_42_ aggregation and evaluate the feasibility of high-field SSNMR for characterizing these heterogeneous assemblies. Unlabeled and uniformly-^13^C,^15^N-labeled Aβ_42_ peptides were expressed recombinantly and purified as described in the Methods section (Fig.2). GD3-Aβ aggregates were prepared under controlled conditions and characterized using complementary biophysical techniques. Thioflavin T (ThT) fluorescence assays were employed to monitor β-sheet formation and aggregation kinetics. Transmission electron microscopy (TEM) imaging revealed morphologically heterogeneous aggregates in the presence of GD3 (Fig.3), consistent with lipid-modulated assembly pathways. To probe the structural features of these assemblies, SSNMR experiments were conducted at both 600 MHz and 1.1 GHz magnetic field strengths on GD3-associated aggregates containing uniformly ^13^C,^15^N-labeled Aβ_42_. Comparative analysis of the spectra demonstrates that ultrahigh-field (1.1 GHz) SSNMR provides substantial improvements in sensitivity and spectral resolution relative to 600 MHz measurements. Notably, resonances corresponding to the C-terminal region of Aβ_42_ exhibit enhanced intensity and resolution at 1.1 GHz, indicating the presence of a relatively well-ordered structural core despite the overall heterogeneity of the sample. These results highlight the advantages of ultrahigh-field SSNMR for studying structurally heterogeneous, lipid-associated amyloid assemblies and provide new insights into GD3-mediated modulation of Aβ_42_ aggregation.

**Figure 2.**
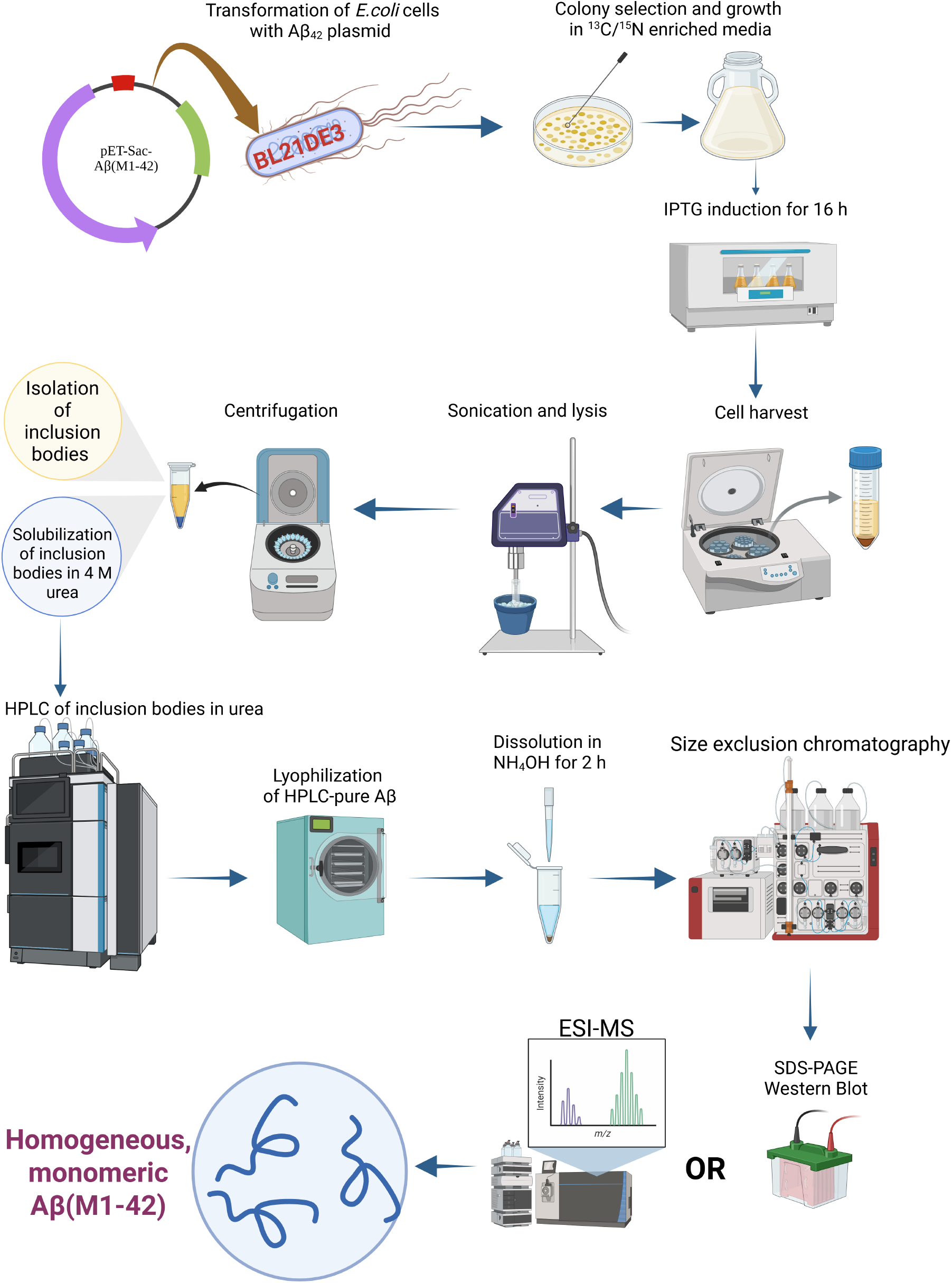
A flow-chart summarizing the steps followed in the biological expression, purification and characterization of monomeric Aβ42.

## Material and Methods

### Materials

Uniformly ^13^C-enriched D-glucose and ^15^N-enriched ammonium chloride were purchased from Cambridge Isotope Laboratories (Tewksbury, MA, USA). The ganglioside GD3 was purchased from Avanti Polar Lipids (Alabaster, AL, USA). *Escherichia coli* BL21(DE3) pLysS Star competent cells were obtained from Invitrogen (Carlsbad, CA, USA).

### Aβ Expression and Purification

Recombinant Aβ(M1-42) was generated in *Escherichia coli* BL21(DE3) pLysS Star expression strains as summarized in Fig.2. For isotopic enrichment, bacterial cultures were propagated in defined minimal media supplemented with ^13^C-enriched glucose and ^15^N-enriched ammonium chloride; non-enriched peptide was produced in standard LB medium. Expression was triggered by the addition of IPTG and maintained for approximately 16 h. Cells were subsequently collected, mechanically disrupted by sonication, and insoluble inclusion bodies were separated. The recovered material was solubilized in 4 M urea, clarified by filtration through a 0.2 μm hydrophilic PVDF membrane, and further purified using high-performance liquid chromatography. Fractions corresponding to Aβ were combined and freeze-dried using a Labconco Triad lyophilizer.

To generate monomeric Aβ_42_, 2–3 mg of the lyophilized peptide was dispersed in 800 μL of ultrapure water and allowed to equilibrate for 30 min. Concentrated ammonium hydroxide was then added to achieve a final concentration of 10–15% (v/v), and the solution was incubated at ambient temperature for 2 h. The sample was subsequently purified by size-exclusion chromatography (SEC) on a Superdex 75 HR 10/30 column preconditioned with 10 mM phosphate buffer (pH 7.4). Separations were carried out at 25 °C with a flow rate of 0.5 mL/min using an ÄKTA FPLC platform (GE Healthcare, Buckinghamshire). Monomeric Aβ eluted in fractions 25-29, and its molecular identity and homogeneity were confirmed by electrospray ionization mass spectrometry.

### Aβ-GD3 aggregate preparation

To generate Aβ-GD3 aggregates, 25 µM of monomeric Aβ, purified via SEC, was incubated with 75 µM of GD3 gangliosides in 10 mM sodium phosphate buffer and 50 mM NaCl at pH 7.4. Reactions were incubated at 37 °C under quiescent conditions for a week. Several reaction batches were set up individually to yield 1mg of total protein and pooled together after incubation. Samples were then lyophilized and resuspended in D_2_O and packed in a 1.6 mm MAS rotor via ultracentrifugation. Samples were characterized by TEM imaging, CD spectroscopy, and SDS-PAGE gel.

### TEM imaging

Samples were prepared by depositing 10 µL of the reaction mixture onto 300-mesh Formvar/carbon-coated copper grids (Millipore Sigma, Burlington, MA, USA; catalog #TEM– FCF300CU) and allowing them to dry for 10 min. The grids were then treated with 10 µL of Uranyless stain (EMS, Hatfield, MA, USA; catalog #22409), blotted to remove excess, and air-dried for an additional 10 min. Imaging was carried out on an HT7800 Hitachi TEM (Minato-ku, Tokyo, Japan) operated at 100 kV. For each sample, images were obtained from at least three grid regions at magnifications ranging from 15,000× to 30,000×.

### NMR experiments

Solid-state NMR spectra were collected at NHMFL on Bruker 600 MHz NMR spectrometer using a 3.2 mm HCN MAS probe, and also at NMRFAM on a Bruker NEO 1.1 GHz spectrometer using a Black Fox (Tallahassee, FL) triple resonance probe in HCN mode. The probe featured a dual-coil design, with an inner solenoid tuned to ^13^C/^15^N and an outer low-electric-field ^1^H resonator. Additional spectra were collected at 600 MHz using an Avance III HD console equipped with a 1.6 mm Phoenix NMR probe of similar coil design. All experiments were performed at 25 kHz with the variable temperature (VT) set to -15 °C. Spectra were referenced to DSS using adamantane as a secondary external standard (downfield ^13^C signal set to 40.48 ppm). 2D ^13^C-^13^C correlation spectra were acquired using 100 ms CORD mixing. 1D Cross-polarization (CP) spectra were obtained with 256 scans, a recycle delay of 1.5 s, a CP contact time of 1 ms, and 100 kHz SPINAL-64 decoupling. 2D spectra were collected with 512 and 1024 indirect points for the 600 MHz and 1.1 GHz instruments, respectively.

## Results and Discussion

### TEM images show the heterogeneity of Aβ_42_-GD3 aggregates

To characterize the assemblies formed in the presence of GD3, transmission electron microscopy (TEM) was employed to examine aggregate morphology. As shown in Fig. 3, the TEM images reveal a heterogeneous population of species, including fibrillar structures along with additional non-fibrillar aggregates. The coexistence of multiple morphologies indicates that GD3 modulates the aggregation pathway of Aβ_42_ and promotes structurally diverse assemblies. These samples were subsequently used for the solid-state NMR experiments described below to investigate their structural features at the molecular level.

**Figure 3.**
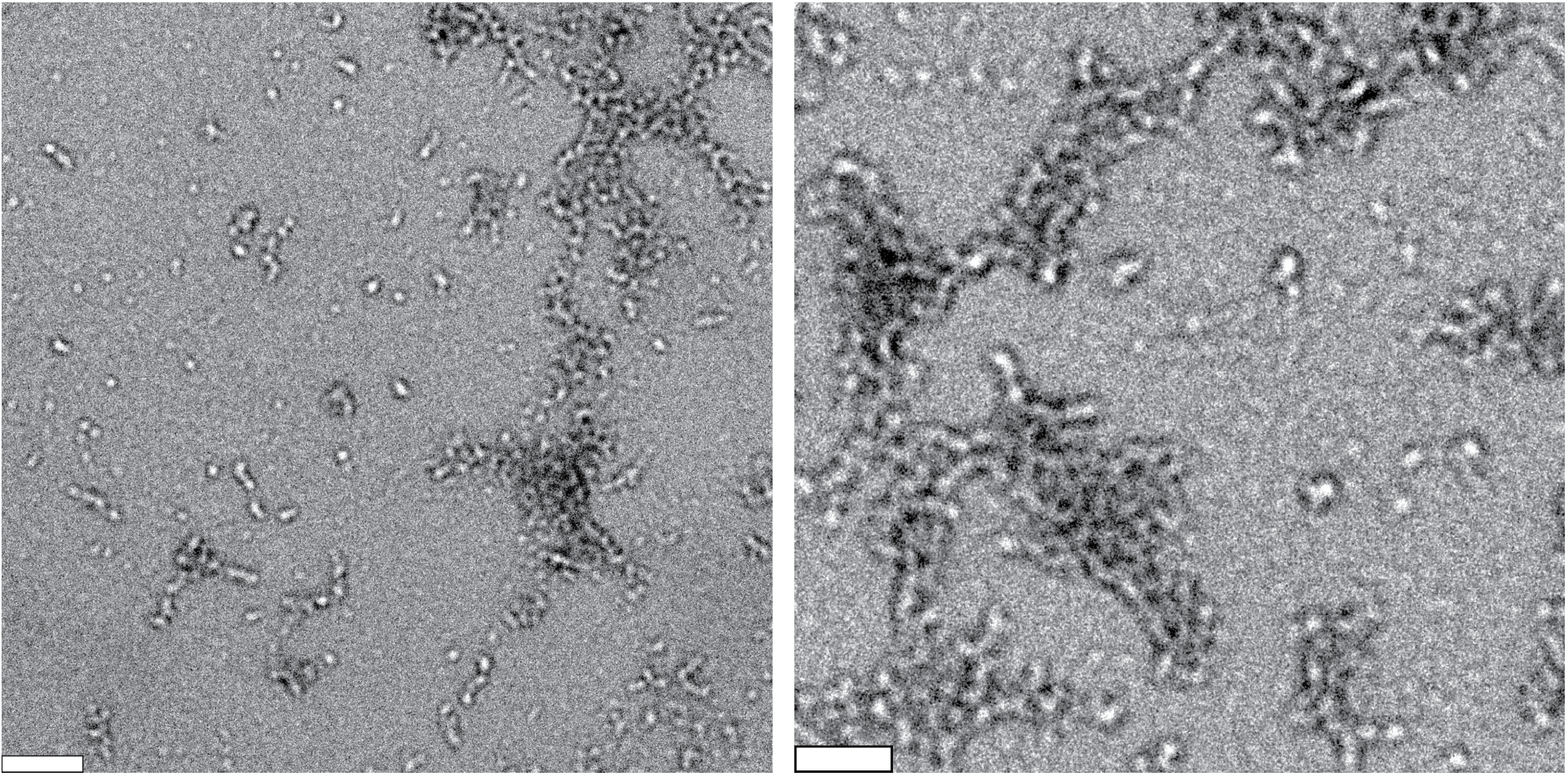
Transmission electron microscopy (TEM) images of Aβ_42_-GD3 aggregates acquired at 15x (left) and 30x (right) magnification. Samples were prepared by incubating 25 μM Aβ_42_ with 75 μM GD3 in 50 mM NaCl and 10 mM sodium phosphate buffer (pH 7.4) under quiescent conditions at 37 °C for one week. Scale bars correspond to 200 nm in (a) and 70 nm in (b).

### Solid state NMR analysis of Aβ_42_–GD3 aggregates

^13^C CPMAS NMR spectra for the Aβ_42_-GD3 aggregates were recorded using a 25 kHz MAS rate, a recycle delay of 1.5 s, and 256 scans (Fig. 4). Under these conditions, the comparison between datasets collected at 600 MHz and 1.1 GHz revealed modest yet noticeable enhancements in signal intensity within the aliphatic regions (10-20 and 40-50 ppm) at the higher magnetic field. These improvements primarily reflect better sensitivity and slightly sharper features at 1.1 GHz; however, the overall cross-polarization efficiency remained largely unchanged. Collectively, these observations indicate that while ultrahigh-field measurements provide incremental gains in spectral clarity for select aliphatic resonances, they do not substantially alter the overall quality of the 1D CPMAS spectra for these heterogeneous Aβ_42_-GD3 assemblies.

**Figure 4.**
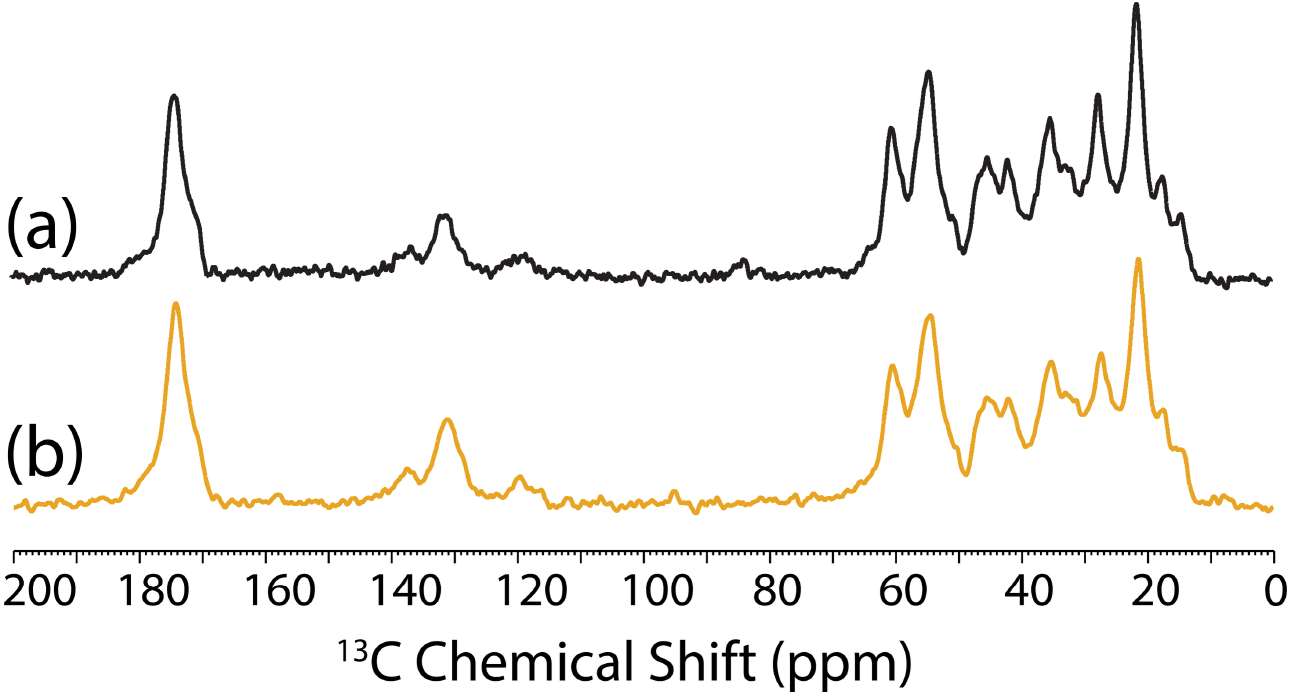
^13^C CPMAS spectra of Aβ-GD3 aggregates acquired at (a) 1.1 GHz and (b) 600 MHz. Measurements were performed using a 1.6 mm MAS probe, at an MAS frequency of 25 kHz and a temperature of -15 °C, with approximately 1 mg of aggregated sample. All NMR acquisition parameters are included in the materials and methods section.

2D ^13^C-^13^C correlation (CORD) spectra acquired with a 100 ms CORD mixing time at 600 MHz and 1.1 GHz reveal prominent intra residue cross peaks for multiple C-terminal residues-Ala30, Ile31, Ile32, Leu34, Met35, and Val40 including characteristic Cα-Cβ correlations, as well as the Leu34 Cβ-Cγ2 correlation (Figs. 5, 6, S1, and S2). These assignments are complemented by clear inter residue contacts, notably Cα-Lys28 - Cα-Gly29 and Cα-Val36 - Cε-Met35 (Fig. 5). Additionally, partially overlapping cross peaks consistent with intra residue contacts are observed within the C-terminal segment for Val36, Val39, Val40 (Cα-Cγ1) and for Ile31, Ile32, Ile41 (Cα- Cγ1 and Cβ-Cγ2), further supporting a tightly packed C-terminal structure in Aβ_42_-GD3 aggregates (Fig. 5). In contrast, signals from much of the central and N-terminal regions are weak or absent in spectra obtained at both 600 MHz and 1.1 GHz, with Arg5 as a notable exception, indicating that these segments are more dynamic and/or less ordered relative to the C-terminus (Fig. 5). This NMR based inference is consistent with our TEM observations, which show poorly ordered Aβ assemblies that lack the well defined fibrillar morphology typically seen for mature Aβ_42_ fibrils; instead, the particles appear protofibrillar, in agreement with prior reports on protofibrillar species of certain Aβ variants^14,15^

**Figure 5.**
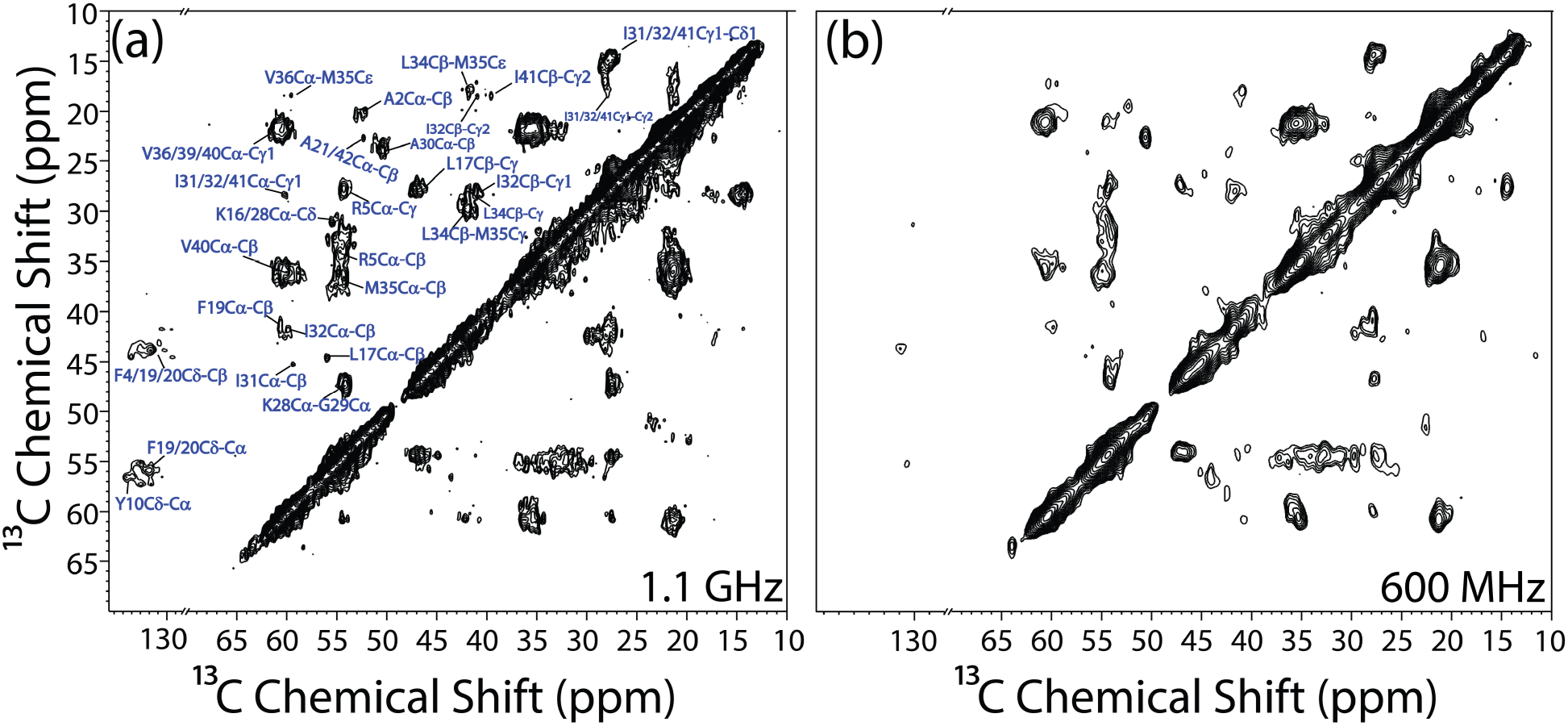
Aliphatic region of 2D ^13^C-^13^C CORD spectra of Aβ_42_-GD3 aggregates acquired at (a) 1.1 GHz and (b) 600 MHz. Measurements were performed using an 1.6 mm HCN probe under 25 kHz MAS. Experimental parameters used include 512 and 1024 t1 increments at 600 MHz and 1.1 GHz, respectively, for the same total t1 eveolution time, 256 scans and a 1.5 recycle delay. Spectra were processed using a sine window function with a sine-bell shift (SSB=2).

**Figure 6.**
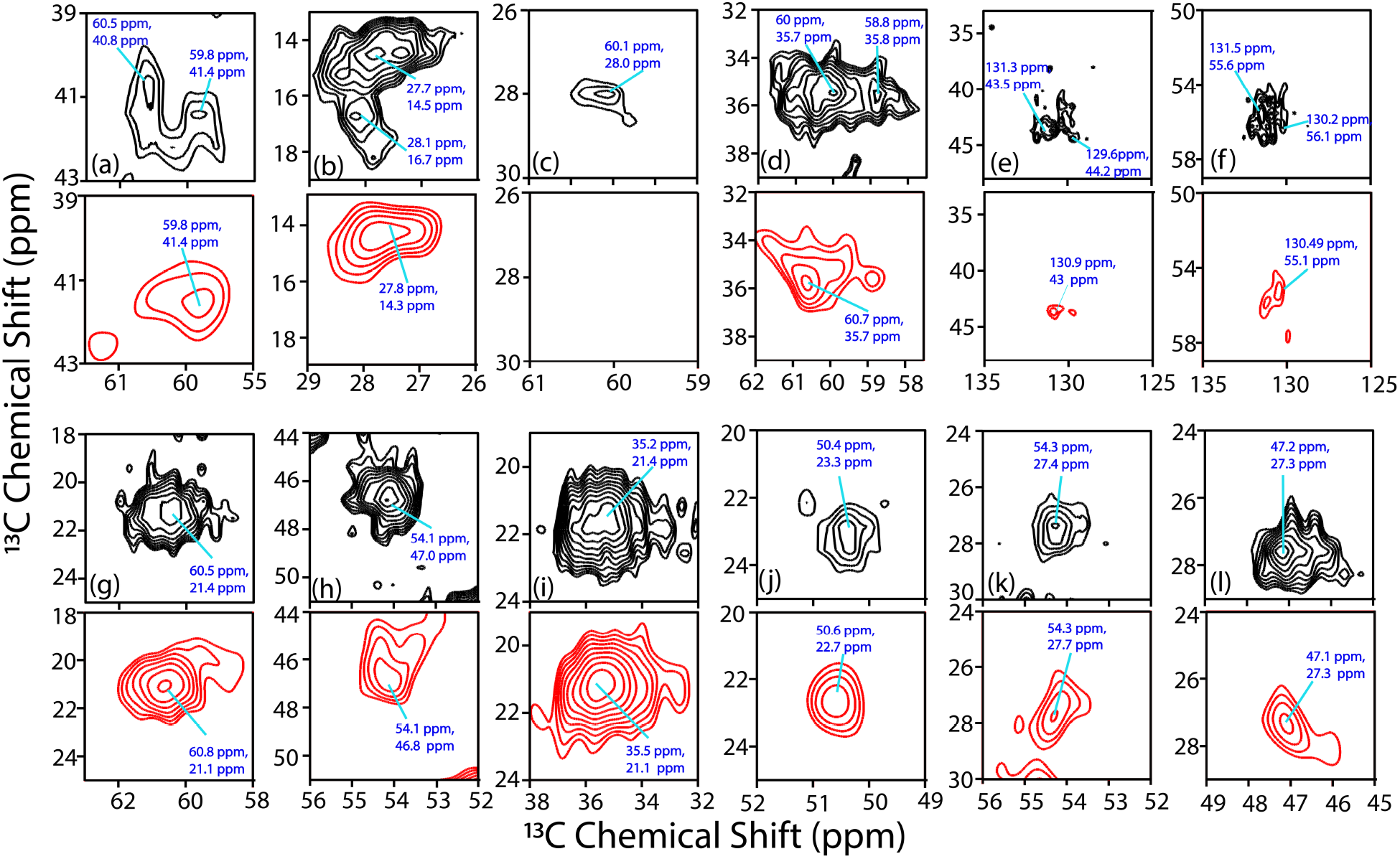
Comparison of the expanded regions of 2D ^13^C-^13^C spectra of Aβ_42_-GD3 (shown in Figure 5) acquired at 1.1 GHz (black) and 600 MHz (red).

A direct comparison of the 600 MHz and 1.1 GHz spectra shows that the ultra-high field provides tangible, but targeted, resolution gains (Figs.6, S1, and S2). At 1.1 GHz, several C-terminal resonances are better resolved, for example, Cβ-Cγ2 and Cα-Cγ of Leu34, and Cβ-Leu34 – Cε-Met35; while weak correlations from the middle region of the peptide emerge, including Leu17 (Cα–Cβ, Cβ–Cγ) and Phe19 (Cα-Cβ) (Fig. 5). These features suggest that experiments carried out at 1.1 GHz can partially reveal signals from relatively heterogeneous segments that otherwise appear as partially broadened at 600 MHz, even if they are not fully ordered (Figs. 5 and 6).

Overall, experiments carried out at 1.1 GHz yield selective improvements in spectral resolution and detectability, the overall spectral quality do not increase dramatically. This behavior underscores reflects the intrinsic feature of heterogeneous assemblies: higher magnetic field strength can enhance spectral resolution and sensitivity, but the dominant determinant of spectral simplification remains the intrinsic orderedness and rigidity of the assemblies formed present in the sample. Consequently, for conformationally heterogeneous or mobile regions, gains from ultra high fields are inherently limited compared with those observed for more rigidly structured amyloid fibrils.^37^

## Conclusions

This study highlights the value of ultrahigh-field solid-state NMR for investigating lipid-modulated, structurally heterogeneous amyloid assemblies. Using Alzheimer’s disease related Aβ_42_ aggregates formed in the presence of the ganglioside GD3, we compared the performance of MAS SSNMR at 1.1 GHz and 600 MHz. Despite the intrinsic heterogeneity and spectral broadening characteristic of lipid-associated aggregates, the 1.1 GHz measurements provided substantial gains in sensitivity and spectral resolution. In particular, 2D ^13^C–^13^C chemical shift correlation spectra acquired using CORD mixing under MAS revealed clearly resolved resonances corresponding to selectively ordered regions of the peptide. The improved spectral quality obtained at ultrahigh field allowed detection of structurally defined segments within the otherwise heterogeneous assemblies, supporting the presence of an ordered fibrillar core in GD3-associated Aβ_42_ aggregates. These observations indicate that even when amyloid samples lack the homogeneity typically required for atomic-resolution SSNMR, experiments performed at ultrahigh-field can recover meaningful structural information. Collectively, this capability opens new opportunities to examine structurally heterogeneous aggregates that are likely more representative of biologically relevant and potentially toxic species, thereby advancing atomic-level understanding of lipid–amyloid interactions in neurodegenerative disease.

## Acknowledgements

This study was supported by NIH (DK132214 to A.R.) and NSF (DMR-2128556) and FSU.

