## Supporting Information for "Higher Magnetic Field NMR Renders Resolution Enhancement on Ganglioside GD3 Catalyzed Aβ_42_ Aggregates"

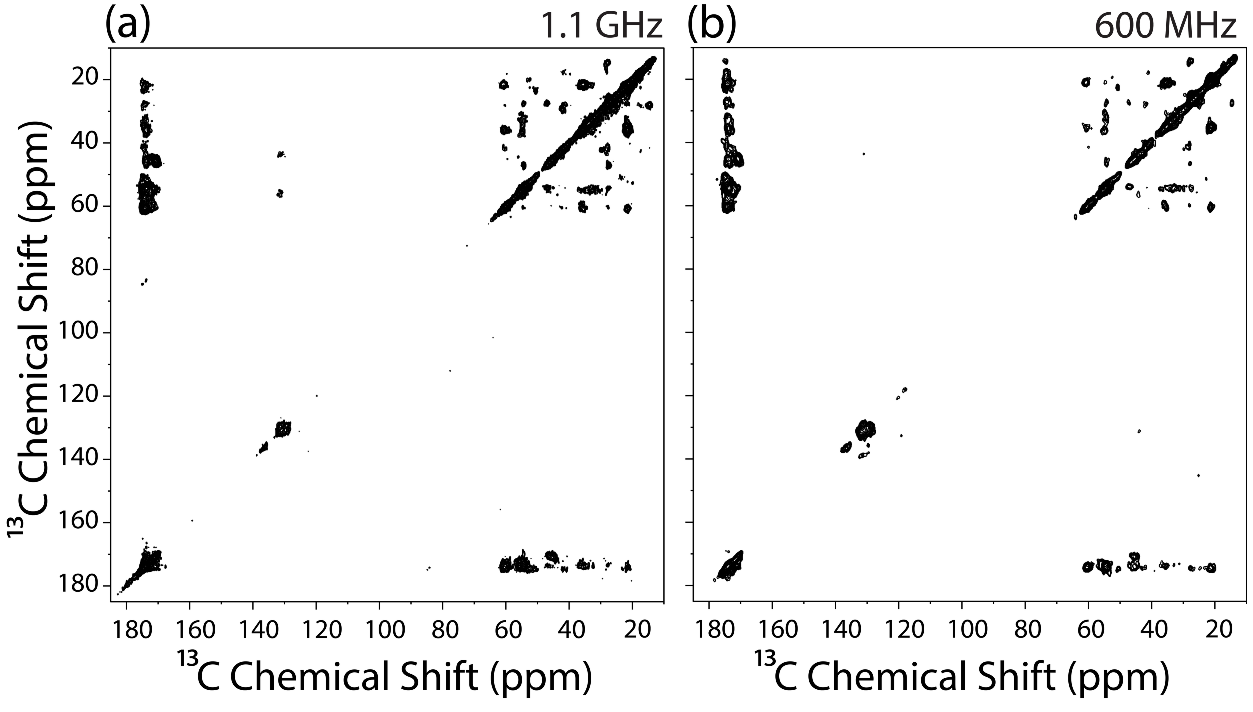


**Figure S1.** 2D $^{13}C$-$^{13}C$ CORD spectra of A$\beta_{42}$-GD3 aggregates acquired at (a) 1.1 GHz and (b) 600 MHz. Expanded regions are shown in Figures 5 and 6.

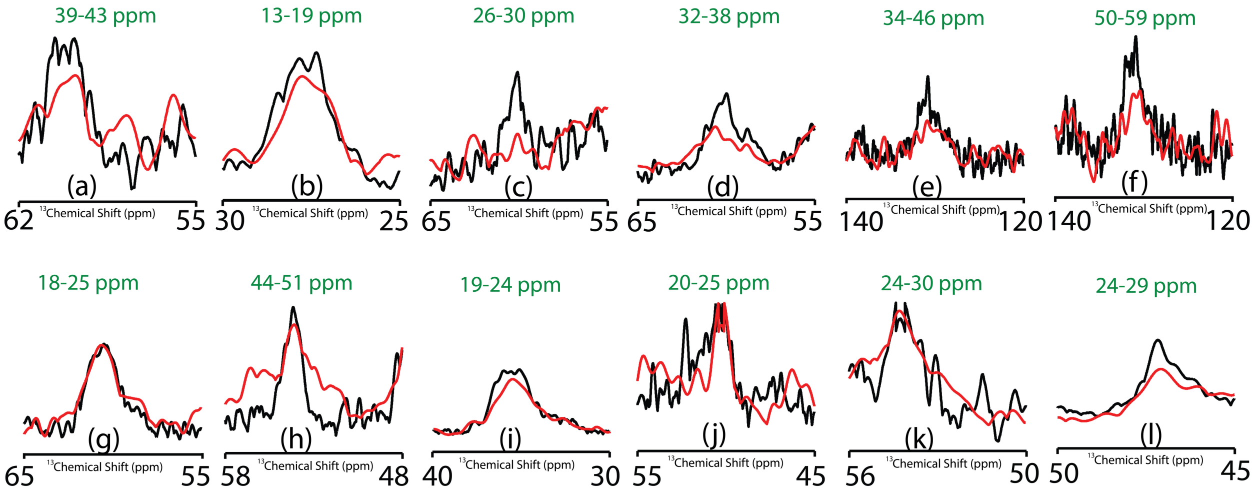


**Figure S2.** 1D projection corresponding to the expanded 2D 13C-13C spectral regions shown in Figure 6. Projection ppm values along the F1 dimensions are indicated in green.
